# Copper-controlled gene expression via transmembrane-induced ribosomal stalling

**DOI:** 10.1101/2025.02.05.636593

**Authors:** Yavuz Öztürk, Kai-Wei Shen, Philip Emmanuel, Fevzi Daldal, Hans-Georg Koch

## Abstract

Regulated gene expression in response to metabolite sensing is a fundamental process for cellular adaptation and survival. Cells have developed diverse strategies to detect and respond to various metabolites in their environment. Here, we have identified a post-transcriptional mechanism in *Rhodobacter capsulatus* that integrates the periplasmic Cu concentration into the translational control of the copper detoxifying enzyme CutO. This is achieved through Cu-induced stalling of the nascent CutF protein inside the ribosomal peptide tunnel during co-translational secretion. Stalling at a C-terminal proline-rich motif overrides the function of elongation factor P (EF-P) and allows melting of an mRNA stem-loop that shields the *cutO* ribosome-binding site. Thus, CutF acts as a transmembrane Cu sensor that controls CutO production via ribosomal stalling. Considering that CutF is a member of the widely distributed bacterial DUF2946 protein family, the mechanism identified here likely represents a conserved bacterial strategy for adapting to toxic heavy metals.

## Introduction

Bacteria can adapt to diverse and rapidly changing environments by regulating gene expression at multiple levels. In particular, multiple mechanisms of translation regulation, *i.e.* regulating translation initiation, elongation, or termination (Njenga *et al*., 2023, Sharma & Woodson, 2020, Starosta *et al*., 2014), complement the well-studied strategies for transcriptional regulation (González-Flecha & Demple, 1997, Gottesman, 2019, Gualerzi *et al*., 2003). This is exemplified by alternative translation initiation mechanisms of leaderless mRNAs (lmRNAs), which lack the canonical ribosome binding site and often encode proteins critical for stress adaptation and survival (Landwehr *et al*., 2021, Beck & Moll, 2018, Gualerzi & Pon, 2015, Cheng-Guang & Gualerzi, 2020).

Another mechanism involves proline-rich ribosome arrest peptides (RAPs), which have been identified in a wide variety of organisms and control translation elongation in response to specific cellular or environmental cues (Murakami *et al*., 2004, Woolstenhulme *et al*., 2013). RAPs cause sequence-dependent translational stalling of nascent polypeptides when they pass through the ribosomal peptide tunnel (Ito & Chiba, 2013, Ito *et al*., 2018). Stalling can regulate the expression of downstream genes by preventing Rho-dependent transcription termination and mRNA degradation, or by increasing the accessibility of the ribosome-binding site of the succeeding gene (Ito & Chiba, 2013). One of the best-characterized examples of a RAP-containing protein is *Escherichia (E.) coli* SecM, which acts as a secretion monitor and senses the translocation activity of the SecYEG translocon (Nakatogawa *et al*., 2004, Oswald *et al*., 2021). SecM is co-translationally secreted via the SecYEG-translocon, however, when the pulling force executed by the SecYEG translocon is insufficient, the C-terminal RAP of SecM gets stalled inside the ribosomal tunnel (Bhushan *et al*., 2011, Butkus *et al*., 2003, Gumbart *et al*., 2012, Niesen *et al*., 2018). This stalling promotes melting of an mRNA stem-loop structure that shields the ribosome-binding site of the down-stream encoded ATPase SecA. Consequently, the increased translation of the secretion motor SecA boosts protein translocation across the SecYEG-translocon (Nakatogawa & Ito, 2002). RAP-regulated synthesis of proteins involved in protein transport appears to be a common strategy for adapting protein secretion to changing conditions and has been experimentally verified for the RAP-containing proteins VemP and MifM as well (Miyazaki *et al*., 2020, Chiba & Ito, 2015). RAP families associated with the protein localization machinery have been identified in multiple bacterial species and many of them contain a RAPP amino acid sequence motif near their respective C-termini (Ishii *et al*., 2015, Morici *et al*., 2024). Although ribosomal stalling is an intrinsic property of arrest peptides present in SecM or VemP, other RAPs, such as TnaC or ErmCL induce stalling only in the presence of their particular substrates tryptophan or erythromycin, respectively. In this respect, they act as metabolic sensors for intracellular metabolites (Ito & Chiba, 2013, Arenz *et al*., 2014, Ramu *et al*., 2011).

While RAP-induced ribosomal stalling upon impaired protein translocation or intracellular metabolite accumulation is well established, the importance of RAPs for sensing extracellular cues is only beginning to emerge. We and others recently identified the RAP-containing DUF2946 protein family, which is involved in copper (Cu) and metal ion homeostasis (Öztürk *et al*., 2021, Öztürk *et al*., 2023, Roy *et al*., 2022). Prototypes of this family are CruR from *Bordetella (B.) pertussis* and CutF from *Rhodobacter (R.) capsulatus* (Roy *et al*., 2022, Öztürk *et al*., 2023, Öztürk *et al*., 2021, Selamoglu *et al*., 2020). Cu binding to CruR alleviates translational arrest and triggers Rho-dependent transcription termination, which in turn prevents production of the downstream-encoded TonB-like Cu-transporter BfrG (Roy *et al*., 2022).

CutF is encoded with the periplasmic multi-copper oxidase CutO and the putative Cu chaperone CutG in the *cutFOG* operon (Öztürk *et al*., 2021). Within the *cutFOG* mRNA, *cutF* and *cutO* are separated by a stem-loop sequence that shields the ribosome binding site of *cutO* and it was suggested that Cu-induced ribosomal stalling of CutF allows unfolding of the stem-loop sequence and CutO synthesis (Öztürk *et al*., 2023). Such a mechanism would be in line with CutO’s role in oxidizing Cu(I) to the less toxic Cu(II) (Andrei *et al*., 2020, Wiethaus *et al*., 2006). Importantly, CutF is dispensable when the stem-loop is mutated (Öztürk *et al*., 2023).

Our bioinformatic analyses revealed that the large majority of DUF2946 proteins contain a Sec signal peptide, a metal binding site, consisting mostly of Cys- and His-residues, and a putative C-terminal arrest peptide with a proline repeat (Öztürk *et al*., 2023). Deleting the signal peptide of CutF, or mutating the Cu-binding or proline-rich motifs resulted in the loss of Cu-dependent CutO production, highlighting their importance (Öztürk *et al*., 2023, Öztürk *et al*., 2021).

Although CruR and CutF belong to the same DUF2946 protein family, there are some striking differences: while Cu binding to CruR is suggested to relieve stalling (Roy *et al*., 2022), it is proposed to induce stalling of CutF (Öztürk *et al*., 2023). Consequently, in the presence of Cu, production of the TonB-like Cu transporter BfrG is prevented while the production of CutO is stimulated. Another difference is the requirement for the signal sequence, which is dispensable in CruR (Roy *et al*., 2022), but essential in CutF (Öztürk *et al*., 2023). Thus, while Cu-dependent regulation of BfrG synthesis is independent of the localization of CruR, the Cu-binding motif of CutF needs to be located in the periplasm for Cu-dependent CutO production. Finally, although consecutive proline residues are poor substrates for peptide bond formation and require a specialized <u>e</u>longation factor <u>P</u> (EF-P) to evade ribosome stalling (Doerfel & Rodnina, 2013, Doerfel *et al*., 2013, Ude *et al*., 2013, Peil *et al*., 2013), EF-P is not involved in relieving CruR stalling in the presence of Cu (Roy *et al*., 2022).

Thus, despite the wide conservation of the DUF2946 protein family in different bacterial species, the mechanisms by which they regulate the production of their associated target gene are still largely unknown. Our data show that nascent CutF senses the periplasmic Cu concentration and that its translation is stalled upon Cu binding. This in turn promotes translation of *cutO* and Cu-dependent CutO production, which is also strictly dependent on EF-P unlike BfrG.

## Materials and Methods

### Bacterial Strains and Growth Conditions

The bacterial strains and plasmids utilized in this study are listed in **Table S1.** *R. capsulatus* strains were cultured under respiratory conditions using either magnesium-calcium peptone yeast extract (MPYE) enriched medium (Daldal *et al*., 1986) or Sistrom’s minimal medium A (Med A) (Sistrom, 1960) at 35 °C. Antibiotics such as kanamycin, gentamycin, or tetracycline were added to the growth media at concentrations of 10, 1, or 2.5 μg/mL, respectively, as required. *E. coli* strains were cultured in lysogeny broth (LB) medium (Bertani, 1951), supplemented with appropriate antibiotics: ampicillin (100 μg/mL), kanamycin (50 μg/mL), tetracycline (12.5 μg/mL) or chloramphenicol (25 μg/ml). For *in vivo* pulse-labeling experiments, the M63 minimal medium containing 18 amino acids (excluding cysteine and methionine), was employed (Pardee & Prestidge, 1959, Jauss *et al*., 2019), with ampicillin added at 50 μg/mL. Constructs harboring the *lac* operator were induced with 1 mM isopropyl β-D-1-thiogalactopyranoside (IPTG) in both rich and minimal media.

### Molecular Genetic Techniques

#### Construction of p*efp*, p*efp::Gm* plasmids and chromosomal inactivation of *efp* in wild-type and Δ*cutFO* strains of *R. capsulatus*

The p*efp* plasmid was constructed by ligating the *efp* gene including its 500 bp upstream (promoter) and 300 bp downstream (transcriptional terminator) regions into the linearized broad host range pRK415 vector **(Table S1)** by KpnI/XbaI digestion. The *efp* gene was amplified by using the EFP-F and EFP-R primers (**Table S2**) and the genomic DNA of the wild-type *R. capsulatus* MT1131 strain as a template. The PCR primers used for the NEBuilder^R^ HiFi assembly method (NEB Lab, USA) contained 20 bp long 5’ overlapping regions between the vector arms and the gene fragments. The quantity and quality of the amplified DNA fragments were checked spectrophotometrically and via agarose gel electrophoresis, respectively, and used in the assembly reactions. HiFi assembly master mix (NEBuilder^R^) was used to assemble the PCR amplified fragment into the linearized conjugative plasmid pRK415 according to the supplier’s protocol. In each case, the total amount of DNA fragments used was ∼ 0.4 - 0.5 pmoles, and the vector to insert ratio was 1:2. The samples were incubated in a thermocycler at 50 °C for 60 minutes, and 4 μL of the assembly reaction was transformed into chemically competent *E. coli* HB101 cells. The isolated plasmids from selected Tet^R^-colonies were confirmed by sequencing.

The Δ(*efp*::*Gm*) deletion-insertion allele in the pRK415 plasmid was obtained by combining a120 bp-long 5’-fragment of the *efp* gene, the gentamicin resistance (Gm^R^) cassette (Prentki & Krisch, 1984), and a 200 bp-long 3’-fragment of the *efp* gene by the HiFi Gibson assembly method, as described by the supplier (NEB lab, MA). The 120 bp-long 5’- fragment and 200 bp 3’-fragment of the *efp* gene were amplified by PCR using the wild type (wt) MT1131 genomic DNA as a template. The PCR primers contained 20 base pairs long 5’ overlapping regions between the vector arms, the gene fragments and the Gm^R^ cassette. The 1.2 kb long Gm^R^ cassette was amplified by using the Gm-F/Gm-R primer pair and pCHB::Gm plasmid as a template (**Table S1**), and the PCR product thus obtained was digested with DpnI to remove the template. HiFi assembly master mix (NEBuilder^R^) was used in a single-step reaction to assemble the N-terminal portion of the target gene, the Gm^R^ cassette, the C-terminal portion of the target gene, and the linearized plasmid pRK415 following the supplier’s protocol as described above. 4 μL of the assembly reaction was transformed into chemically competent *E. coli* HB101 cells. After sequencing, the correct clone in the plasmid pRK415 was conjugated into the GTA overproducer *R. capsulatus* Y262 strain (Yen *et al*., 1979) via triparental crosses (Koch *et al*., 1998) (**Table S1**). Y262 strain carrying the Δ(*efp*::*Gm*) allele in pRK415 plasmid was grown on MPYE medium under the photosynthetic condition for 3-4 days to reach the late stationary phase to enrich the GTA particles. The GTA particles were then isolated by filtering aseptically through a 0.45 μ, and then 0.2 μ disposable filters, and used to inactivate the chromosomal copy of the desired genes in wild-type *R. capsulatus* MT1131 and Δ*cutFO* strains selecting for Gm^R^ (Koch *et al*., 1998) (**Table S1**).

### Construction of CutF variants

The membrane-bound CutF (CutF_TM_) variant was constructed by mutating the signal peptide cleavage site of CutF (VA_28_A_29_ to VI_28_I_29_) in the p*cutFOG* plasmid. The CutF(EP)- F/CutF(SPm)-R and CutF(SPm)-F/CutTer-R primer pairs were used to amplify the *cutFOG* fragment carrying the VI_28_I_29_ mutation and 20 bp overlapping sequences. The amplified fragments were subjected to *Dpn*I digestion to remove the template DNA (p*cutFOG*) and purified by the Qiagen PCR purification kit (Qiagen, Hilden, Germany). Fragments were integrated into the linearized pRK415 plasmid by NEBuilder^R^ HiFi assembly cloning method (NEB Lab, United States) as described above.

p*cutF*_P-A_*OG* carrying PP_115_P to PA_115_P substitution of CutF in p*cutFOG* was constructed by using the cutF(EP)-F/CutF-CterM-Rv and CutF(P115A)-F/CutTer-R (**Table S2**) primer pairs and p*cutFOG* plasmid as a template to amplify the mutated fragments. After PCR amplification, amplified products were treated with *Dpn*I digestion to remove the template plasmid, and purified by the Qiagen PCR purification kit (Qiagen, Hilden, Germany). Fragments containing the 20 bp overlapping sequences were assembled into the linearized (KpnI/XbaI digested) pRK415 plasmid by NEBuilder^R^ HiFi assembly cloning method (NEB Lab, United States). 4 μL of the assembly reaction were transformed into chemically competent *E. coli* HB101 cells. The isolated plasmids from selected Tet^R^ colonies were confirmed by sequencing.

For site-directed cross-linking experiments, amber stop codons were inserted into pRS1-CutF, encoding *cutF* under the control of the T7 promoter (Öztürk *et al*., 2021). For the construction of S93, L95, P97, A99, P101, L104 andA105 amber stop codon substitutions of CutF, the Q5 mutagenesis Kit (NEB Lab, MA) was used with the mutagenic primer CutF(S93TAG)-F, CutF(L95TAG)-F, CutF(P97TAG)-F, CutF(A99TAG)-F, CutF(P101TAG)-F, CutF(L104TAG)-F, CutF(A105TAG)-F, and CutF(AmberTGA)-R **(Table S2)** and 10 ng of pRS1-CutF as a template. 5 μL of the KLD reaction mixture (NEB Lab, United States) was transformed to chemically competent *E. coli* NEB® 5-alpha cells. The presence of the amber stop codon was confirmed by DNA sequencing, and the plasmids were subsequently transformed into *E. coli* MC4100 via TSB transformation (Chung & Miller, 1988).

The pRS*cutFO* and its variants carrying the stem-loop mutation (*cutF*_SLm_*O*), the PP_115_P to PA_115_P substitution (*cutF*_P-AP_*O*), or their combinations (*cutF*_P-AP&SLm_*O*) were constructed by using the NEBuilder^R^ HiFi assembly cloning method (NEB Lab, United States) with pRS1cutF-F/ pRScutO-R primer pairs for pRS*cutFO*, pRScutF(P-A)O-F/ pRScutF(P-A)O-R primer pairs for pRS*cutF*_P-AP_*O and CutO-SL-F/ CutO-SL-R* primer pairs for pRS*cutF*_SLm_*O* (**Table S2**). The PCR products were digested with DpnI to remove the template DNA and purified by a Qiagen PCR purification kit (Qiagen, Hilden, Germany). The NEBuilder^R^ HiFi assembly cloning method (NEB Lab, United States) and transformation procedures like those described above were used. The resulting plasmid pRSLepB-CutF(fusion) was confirmed by DNA sequencing and transformed into *E. coli* MC4100 via TSB transformation.

### Biochemical and Biophysical Techniques

#### Preparation of the Periplasmic Fraction for Multicopper Oxidase Activity Assays and SDS-PAGE Analysis

The multicopper oxidase (CutO) containing periplasmic fraction of *R. capsulatus* was isolated from 10 or 50 mL cultures grown overnight (∼20 hours) in MPYE medium under respiratory conditions at 35°C with 110 rpm shaking, in the presence or absence of 10 μM CuSO_4_ supplementation (Trasnea *et al*., 2016a, Trasnea *et al*., 2016b). Cells were harvested and washed at 4 °C with 12 mL of 50 mM Tris-HCl (pH 8.0), and the resulting pellet was resuspended to a final concentration of 10 mL per gram of cell wet weight in SET buffer (0.5 M sucrose, 1.3 mM EDTA, 50 mM Tris-HCl, pH 8.0). The suspension was incubated with 600 μg/mL lysozyme at 30 °C for 60 minutes. Following incubation, the suspension was centrifuged at 13,000 rpm for 30 minutes at 4°C, and the supernatant (*i.e.* the periplasmic fraction) was collected. This periplasmic extract was either used immediately for multicopper oxidase activity assays and SDS-PAGE, or stored at -80 °C.

### Multicopper Oxidase Activity Assay

The oxidation of 2 mM of 2,6-dimethoxyphenol (2,6-DMP) was assessed in a reaction mixture containing 100 mM sodium acetate buffer (pH 5) and 250 μM CuSOC. The reaction was catalyzed using 50 μg of the periplasmic protein fraction. Absorbance measurements were performed at 468 nm using an Ultrospec 3100 Pro UV-Vis spectrophotometer with endpoint detection. The concentration of oxidized 2,6-DMP was determined using a molar extinction coefficient (ε) of 14,800 MC¹ cmC¹ for oxidized 2,6-DMP (Öztürk *et al*., 2021).

### Immunodetection

Proteins were separated by 12%, 15%, or 4-15% gradient SDS-PAGE electrophoresis. Following electrophoresis, proteins were transferred onto nitrocellulose or PVDF Immobilon-P membranes (GE Healthcare, Germany). Anti-Flag tag and Anti-Bacterial Alkaline Phosphatase (BAP, PhoA) antibodies were obtained from Sigma or Millipore (Temecula, USA). Peroxidase-coupled anti-mouse IgGs from SeraCare (medac GmbH, Wedel, Germany) were used as secondary antibodies with ECL (GE Healthcare). Chemiluminescence was detected by the FUSION FX or FUSION FX 7 edge imaging system (Vilber Lourmat).

### *In vivo* pulse labeling for ribosome-associated polypeptide chains of LepB–CutF and CutF

A 3 mL pre-culture of *E. coli* BW25113 (wt) and *E. coli* Δ*efp* strains harboring the pRS1LepB–CutF or pRS1-CutF plasmid were grown on LB medium containing 100 μg/mL ampicillin The cultures were incubated overnight at 37 °C with 800 rpm shaking. Subsequently, 2 mL of each culture were harvested, washed twice with sterile media, and resuspended in 200 μL of M63 minimal medium containing 18 amino acids (M63 18-aa). A total of 150 μL of these cell suspensions was then used to inoculate 10 mL of fresh M63 medium supplemented with 25 μg/mL ampicillin. Cultures were incubated at 37 °C with shaking at 180 rpm until an optical density (OD600) of 0.5–0.8 was reached. At this stage, approximately 2 × 10C cells from each culture were collected, and transferred into 2 mL Eppendorf tubes, and the volume in each tube was adjusted to 2 mL with fresh M63 medium. To inhibit endogenous *E. coli* RNA polymerase, 50 μg/mL of rifampicin was added to the samples, followed by incubation at 37 °C for 15 minutes. The production of LepB–CutF variants was induced by the addition of 0.1 M IPTG along with 2 μL of L-[^35^S] methionine-cysteine mix (7 mCi/mL; PerkinElmer Life Sciences) (Jauss *et al*., 2019). Samples of 100 μL were collected at 1-, 2-, 3-, and 6-minutes post-induction and immediately precipitated with 10% trichloroacetic acid (TCA), followed by incubation on ice for 30 minutes. Precipitated proteins were recovered by centrifugation at 13,500 rpm for 15 minutes at 4 °C. The resulting protein pellets were denatured by resuspension in 25 μL of SDS sample buffer and incubated at 56 °C with continuous shaking at 1,400 rpm for 15 minutes. Proteins were then resolved by 15% SDS-PAGE. The gel was subsequently dried and analyzed by phosphorimaging.

### Isolation of a translation-competent S135 fraction

For isolating a translation-competent S135 extract, *E. coli* C43 pEVOL and *E. coli* Δ*efp* pEVOL cells were cultured in an S130 medium (0.8 g/L yeast extract, 9 g/L tryptone-peptone, 5.6 g/L NaCl, 1 mM NaOH, and 20% (w/v) glucose) containing chloramphenicol (25 μg/ml) until reaching an optical density (ODCCC) of 1.0–1.2. Cells were harvested by centrifugation at 3,234 x g for 30 minutes at 4 °C. The resulting pellets were resuspended in S30 buffer (10 mM TeaAc, pH 7.5, 60 mM potassium acetate, 14 mM magnesium acetate, and 1 mM dithiothreitol (DTT)). Cell walls were digested with lysozyme in the presence of the protease inhibitor phenylmethylsulfonyl fluoride (PMSF) for 20 min at room temperature. Following digestion, the sample was subjected to sonication at 23% amplitude for 7 x, with 15-second pulses followed by 45-second pauses by Ultrasonic Homogenizers (Bandelin Sonoplus). Cells were then centrifuged using a Beckman table-top ultracentrifuge at 33,719 x g in the TLA 100.3 rotor for 10 minutes at 4 °C. For *in vitro* translation of the endogenous mRNAs, the supernatant was subsequently incubated in a read-out reaction mixture consisting of 1 mM Tris-acetate (pH 7.5), 1 mM DTT, 1 mM magnesium acetate, 1 mM 18 amino acid mix, 1 mM cysteine and methionine, 250 mM ATP, 200 mM phosphoenolpyruvate (PEP), and 2 mg/mL pyruvate kinase. The reaction mixture was incubated on a Thermo mixer at 450 rpm, 37 °C, for 1 hour. Following incubation, the sample was dialyzed by using a dialysis tube (14,000-16,000 Da) against S30 buffer (10 mM Tris-acetate pH 7.5, 14 mM Mg acetate, 60 mM KOAc, freshly prepared1 mM DTT). S135 fraction containing all essential components for translation was concentrated via ultracentrifugation at 345,287 x g (80,000 rpm) for 9 minutes at 4 °C by using TLA 100.3 rotor.

### Isolation of ribosomes and cytosolic translation factors (CTF)

High-salt washed ribosomes and cytosolic translation factors (CTF) were purified from wild-type and JW4107(Δ*efp*) strains. Cells were cultured in S130 medium until reaching an ODCCC of 1.6–1.8 and were harvested at 5,422 × g for 10 min at 4 °C. Pellets were washed in CTF buffer (50 mM Tris-acetate (pH 7.5), 50 mM potassium acetate, 15 mM magnesium acetate, 1 mM DTT) and lysed using an EmulsiFlex C3. Cell lysate was cleared at 30,000 × g for 30 min at 4 °C using an SS34 rotor, and the supernatant was further separated by ultracentrifugation at 184,000 × g for 2.5 h using a Ti50.2 rotor into supernatant, containing soluble material and pellet containing ribosomes and membranes. The supernatant was further fractionated through a 10–30% sucrose gradient at 39,000 rpm for 22 h in a TH-641 swing-out rotor at 4 °C. Fractions of 500 μL were withdrawn from the gradient and translation competent fractions were pooled as CTF. Translation competency was tested as described below. The ribosomal pellets after ultracentrifugation, as described above were resuspended in high-salt buffer (50 mM Tris-acetate, 50 mM potassium acetate, 15 mM magnesium acetate, 1 mM DTT, pH 7.5), layered on a 50% sucrose cushion, and centrifuged at 344,000 × g for 1 h in a TLA 100.3 rotor. The pellet was resuspended in CTF buffer and centrifuged through a 10-40 % sucrose gradient in high-salt buffer at 29,000 rpm for 17 h in a TH-641 swing-out rotor (Thermo Fisher Scientific). The ribosomal fractions were withdrawn from the gradient, pooled, and concentrated at 85,000 rpm for 1h in a TLA120.3 rotor and resuspended in CTF buffer at pH 7.5 with 1 mM DTT.

### *In vitro* protein synthesis

*In vitro* protein synthesis was performed in a coupled *in vitro* transcription/ translation system in *E. coli* as described before (Öztürk *et al*., 2021, Koch *et al*., 1999). A purified transcription/translation system composed of cytosolic translation factors (CTF) and high salt-washed ribosomes was used. The ^35^S-Methionine/ ^35^S-Cysteine labeling mix was obtained from Hartmann Analytics (Braunschweig, Germany). T7-based expression plasmid pRS1 carrying CutF variants were used as 20 ng/μl final concentration. The samples were incubated at 37 °C for 30 min with gently shaking with 450 rpm. After synthesis, the reaction was mixed with 10% TCA on a 1:1 volume, for 30 min on ice. Precipitated proteins were pelleted in a precooled table-top centrifuge and resuspended in a 30 μl loading dye, which was prepared as described before (Steinberg *et al*., 2020). Resuspended proteins were separated on a 15% SDS-PAGE and visualized by phospho-imaging.

### *In vivo* site-directed cross-linking

p-benzoyl-l-phenylalanine (pBpa) for cross-linking was obtained from Bachem (Bubendorf, Switzerland). The amber-stop codon variants of CutF on a pRS1 were transferred to *E. coli* C43(DE3) pEVOL strain (Panahandeh & Muller, 2010, Ryu & Schultz, 2006) and cultured overnight in 2 ml LB medium at 37 °C with ampicillin and chloramphenicol. 200 μl of the overnight culture were used for inoculation of 10 ml LB medium supplemented with 10 μl pBpa (final concentration 0.5 mM, dissolved in 1 M NaOH). The cultures were further incubated at 37 °C until they reached the early exponential growth phase (OD_600_ = 0.5-0.8) and induced with 1 mM IPTG and 0.02% arabinose. After induction, the cultures were grown for 2 hours at 37 °C, and OD_600_ values were measured for calculating 4x10^8^ cells. The volume required 4x10^8^ cells were exposed to UV light (365nm) on ice for 30 minutes (UV chamber: BLX-365, from Vilber Lourmat). After UV irradiation samples were precipitated with 5% TCA (final concentration) for 30 min on ice. Precipitated proteins were pelleted in a precooled table-top centrifuge and resuspended in a 30 μl loading dye, which was prepared as described before (Steinberg *et al*., 2020). Resuspended proteins were separated on a 15% SDS-PAGE and visualized by phospho-imaging.

### *in vitro* site-directed cross-linking

For *in vitro* crosslinking experiments two approaches were used. In the first approach, ribosomes carrying pBpa at position 71 of the ribosomal protein uL23 (uL23(G71-pBpa)) were isolated as described (Denks *et al*., 2017, Knüpffer *et al*., 2019) and used for coupled *in vitro* transcription/translation of CutF in the CTF system (Koch *et al*., 1999). In the second approach, the S135 fraction (containing ribosomes and amber stop codon-suppressing tRNAs and an engineered tRNA-synthetase) from *E. coli* C34 pEVOL strain was used for the amber stop codon containing CutF variants. The pRS-CutF amber stop codon variants were *in vitro* synthesized using the S135 fraction and in the presence of freshly prepared pBpA (40μM). In both approaches, after 30 min in vitro translation, the reactions (50 μl) were exposed to UV light using a Biolink 365 nM-crosslinking chamber (Vilber-Lourmat) for 30 min on ice for inducing the crosslinking reactions. Samples were then TCA-precipitated, separated on PAGE, and analyzed by phospho-imaging.

## Results

### Elongation factor P is required for CutF-dependent Cu induced CutO production

In a previous study, we demonstrated that the C-terminal proline-rich motif of CutF is essential for Cu regulated CutO production (Öztürk *et al*., 2023, Öztürk *et al*., 2021). This is intriguing because peptide-bond formation between proline residues is intrinsically difficult and depends on the presence of Elongation factor P (EF-P). EF-P is conserved in all domains of life and prevents stalling of proline stretches inside the ribosome (Hummels & Kearns, 2020). However, ribosomal stalling can also be physiologically relevant and has been shown to control gene expression of the ATPase SecA or the SecDF complex (Murakami *et al*., 2004, Ishii *et al*., 2015). For monitoring whether ribosomal stalling of CutF is required for CutO expression, we generated an *R. capsulatus* Δ*efp* strain. Although EF-P was initially considered to be essential, recent data demonstrate that bacterial Δ*efp* strains are viable at low growth rates, which we also observe here for the *R. capsulatus* Δ*efp* strain **(Fig. 1A)**. *R. capsulatus* cells lacking both *cutFO* and *efp* were also viable and their growth defect was restored by providing *efp* in trans **(Fig. 1A)**.

**Figure 1.**
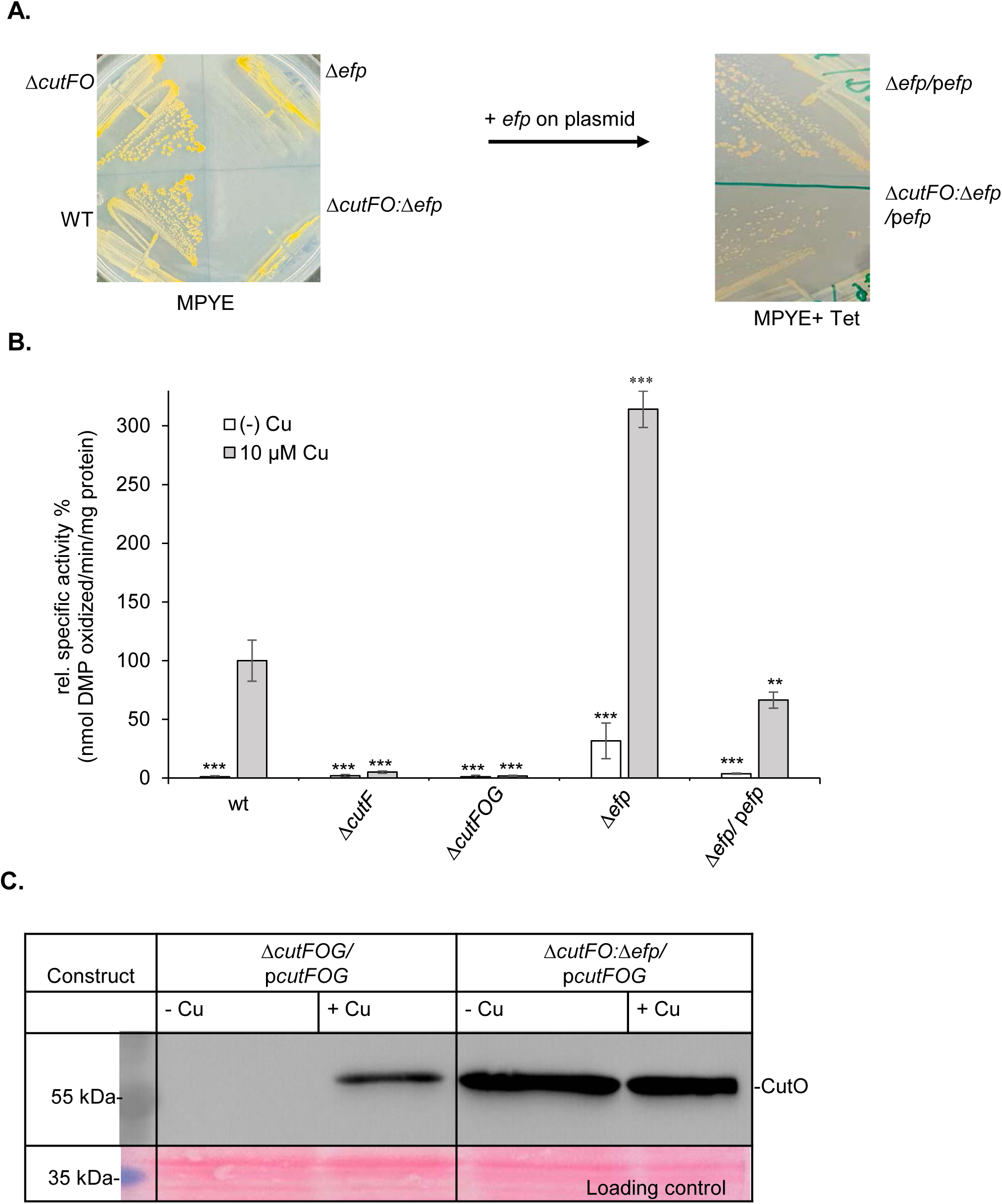
Elongation factor P (EF-P) plays a vital role in Cu-regulated CutO production in *R. capsulatus*. **(A)** Respiratory growth on MPYE medium of different *R. capsulatus* strains. The right image shows the Δ*efp* and Δ*cutFO:*Δ*efp* strains harboring a plasmid encoded *efp* copy. **(B)** CutO activity in different *R. capsulatus* strains. Periplasmic fractions were isolated from the indicated strains, grown in MPYE medium supplemented without or with 10 mM CuSO_4_. 50 mg periplasmic soluble protein was used for the activity measurement using 2,6-dimethoxyphenol oxidation as read-out. The activity of the wild type (wt) grown in the presence of Cu was set to 100% and the relative activities of the indicated strains were calculated. Three independent experiments were performed with three technical replicates and the error bars reflect the standard deviation (n = 9). Statistical analyses were performed with the Satterthwaite corrected two-sided Student t-test, using the activity of the wt + Cu as reference (set to 100%) *, P <0.05; **, P < 0.01; ***, P <0.001. Related to Figure S1. **(C)** Immunoblot analyses of the strains were performed by separating 100 mg of periplasmic soluble protein on 15% SDS PAGE. After transfer to nitrocellulose membrane, α-Flag antibodies were used for the detection of CutO as described in Materials and Methods. The same membrane was stained with Ponceau solution as loading control.

For analyzing whether the absence of EF-P influences the CutO activity, we monitored it in the periplasmic fraction of the Δ*efp* strain via the 2,6-DMP assay. In wild type cells, CutO activity is undetectable in cells grown without Cu supplementation, but is strongly stimulated upon Cu supplementation of the media (**Fig. 1B**). In the Δ*cutF* and Δ*cutFOG* strains no CutO activity was detectable, but the activity was restored in the presence of plasmid-encoded copies of the *cutFOG* operon (p*cutFOG*, **Fig. S1).** In the Δ*efp* strain, high CutO activity was already observed without Cu supplementation (**Fig. 1B**), indicating that absence of EF-P mimics the effect of Cu supplementation on CutO activity. Cu supplementation had a synergistic effect and further increased the CutO activity in the Δ*efp* strain. In the presence of a plasmid-borne *efp* copy, the CutO activity was comparable to the wild type activity, indicating that the absence of EF-P stimulated either the steady-state levels or the activity of CutO **(Fig. 1B)**. For differentiating between the effects of Cu and EF-P on CutO synthesis or activity, the levels of the Flag-tagged CutO were determined by immune detection. In wild type cells, CutO was only detectable when cells were grown on Cu-supplemented media; however, the CutO levels were largely independent of Cu supplementation in the Δ*efp* strain (**Fig. 1C**). Although Cu supplementation does not significantly increase the CutO levels in the Δ*efp* strain, its activity was strongly stimulated, possibly because CutO contains four Cu centers in its active site (Sakuraba *et al*., 2011), and thus, Cu serves not only as CutO substrate, but also as its cofactor.

For validating that Cu also stimulated CutO activity, we deleted *efp* in the Δ*cutFO* background. The Δ*cutFO* strain shows no CutO activity, and this was not influenced by the simultaneous deletion of *efp* (**Fig. 2A**), but was rescued by a plasmid-borne copy of the *cutFOG* operon. In contrast, a plasmid containing CutF with a mutated Cu-binding motif (p*cutF_C-A_OG*) failed to restore CutO activity. Importantly, the Δ*cutFO*:Δ*efp* strain carrying either p*cutFOG* or p*cutF_C-A_OG* showed high CutO activity already in the absence of Cu, which was further stimulated by Cu supplementation **(Fig. 2A)**. Thus, the increased CutO activity upon Cu supplementation when *efp* is deleted is mainly caused by Cu serving as cofactor of CutO. Immune detection confirmed that comparable amounts of CutO were produced in Δ*cutFO*:Δ*efp* strain complemented with either p*cutFOG* or p*cutF_C-A_OG* **(Fig. 2B)**. The data also suggest that the absence of EF-P had an even stronger effect on CutO production than the addition of Cu, because the Cu-induced CutO levels in the absence of EF-P were higher than in its presence (**Fig. 2B**).

**Figure 2.**
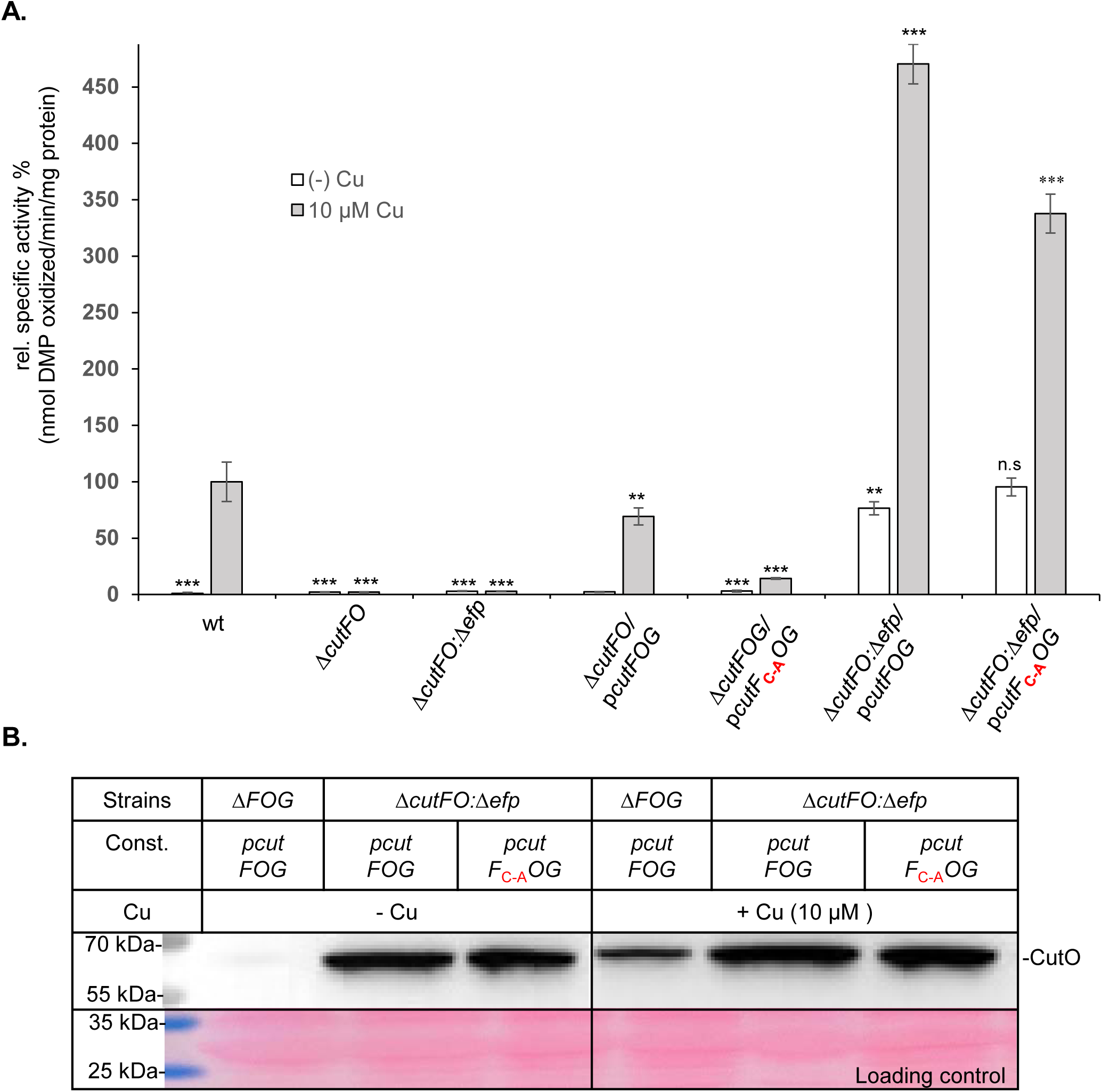
Cu-regulated CutO production is dependent on the presence of both the secreted CxxxC motif and EF-P. **(A)** Periplasmic fractions were isolated from the indicated strains supplemented without or with 10 mM CuSO_4_. The CutO activity was determined by using 50 mg periplasmic soluble total proteins as described in the legend to Fig. 1B. The activity of the wild type (wt) grown in the presence of Cu was set to 100% and the relative activities of the indicated strains were calculated. Three independent experiments were performed with three technical replicates and the error bars reflect the standard deviation (n = 9). Statistical analyses were performed as described in Fig. 1 and n.s. denotes no significant changes. The plasmid pCutF_C-A_OG encodes a CutF variant with a mutated Cu-binding motif. Related to Figures S2 and S3. **(B)** Immunoblot analyses of the strains harboring the wt CutF and CutF_C-A_ variant. 100 mg of periplasmic soluble protein were separated on 15% SDS PAGE and a-Flag antibodies were used for the detection of CutO as described in Materials and Methods. The same membrane was stained with Ponceau solution as loading control.

We had recently shown that a mutation within the mRNA stem-loop that separates *cutF* and *cutO* also allows for Cu-independent CutO production (Öztürk *et al*., 2023) **(Fig. S1A & B**). Thus, Cu-regulated CutO production depends on the presence of the stem-loop that shields the ribosome-binding site of *cutO*. These data support a model in which CutO production depends on CutF stalling, which is induced by either deleting EF-P or by binding Cu to the CxxxC motif of CutF. Moreover, mutation of the CxxxC motif inhibits CutO production, regardless of Cu availability, whereas the Cu-binding motif is dispensable for CutO production when EF-P is deleted. In summary, overall findings demonstrate that control of Cu-induced CutO production is influenced by both EF-P and Cu binding to CutF.

### Disruption of the C-terminal PPP proline repeat abolishes the requirement of EF-P for Cu-regulated CutO production

The specific involvement of EF-P in peptide bond formation of the C-terminal proline-rich motif of CutF (P_109_EPEG<u>PPP</u>RL_118_) was validated by analyzing a variant in which the native PPP motif was replaced by PAP. In the presence of CutF_PAP,_ neither the addition of Cu nor the deletion of EF-P resulted in a significant stimulation of CutO activity (**Fig. 3A**). Immune detection confirmed that Cu-dependent CutO production was not observed when CutF contained the mutated PAP-motif (**Fig. 3B**). Similarly, a strain lacking EF-P (Δ*cutFO:*Δ*efp*) also did not produce CutO irrespective of Cu supplementation when the proline motif of CutF was mutated (**Fig. 3C**). The importance of the proline-rich motif in CutF for Cu-dependent CutO production provides further support for a model in which Cu binding to CutF induces ribosomal stalling that in turn allows for CutO production.

**Figure 3.**
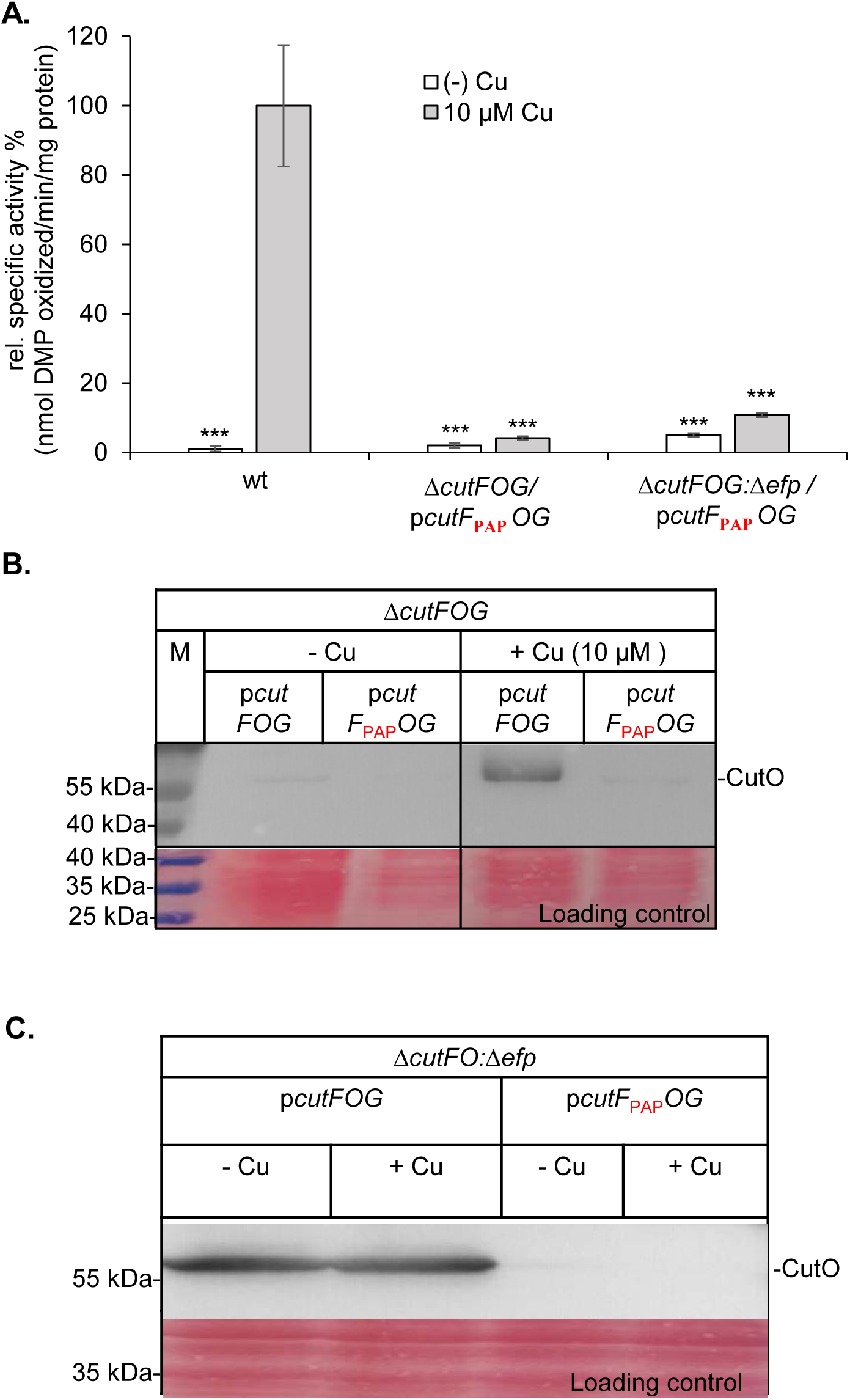
Proline repeats at the C-terminus are required for Cu-regulated CutO production. **(A)** The CutO activity (n = 9) was determined by using 50 mg periplasmic soluble total proteins as described in the legend of Fig. 1B. The plasmid pCutF_PAP_OG encodes a CutF variant with a mutated C-terminal proline motif. **(B) & (C)** Immunoblot analyses of the strains harboring the wt CutF and CutF_PAP_ variant were performed by using 100 mg of periplasmic soluble proteins separated on 15% SDS PAGE and a-Flag antibodies were used for the detection of CutO as described in Materials and Methods. The same membrane was stained with Ponceau solution as loading control.

### CutF is stalled via its proline-rich motif inside of the ribosomal tunnel

We have previously shown that CutF is co-translationally secreted into the periplasm of *R. capsulatus* and rapidly degraded (Öztürk *et al*., 2023, Selamoglu *et al*., 2020), which prevents the reliable detection of stalled CutF-intermediates. For overcoming this hurdle, we heterologously expressed in *E coli* a LepB-CutF reporter construct, which consisted of the first two transmembrane domains of LepB, fused to a signal sequence-less CutF variant containing an additional 23 amino acids at the C-terminus (**Fig. 4A**). Membrane anchoring of CutF does not impair its function as Cu-dependent regulator of CutO synthesis (**Fig. S2**); however, deleting CutF’s signal sequence drastically reduces CutO production and activity both in the absence and presence of EF-P (**Fig. S3**)

**Figure 4.**
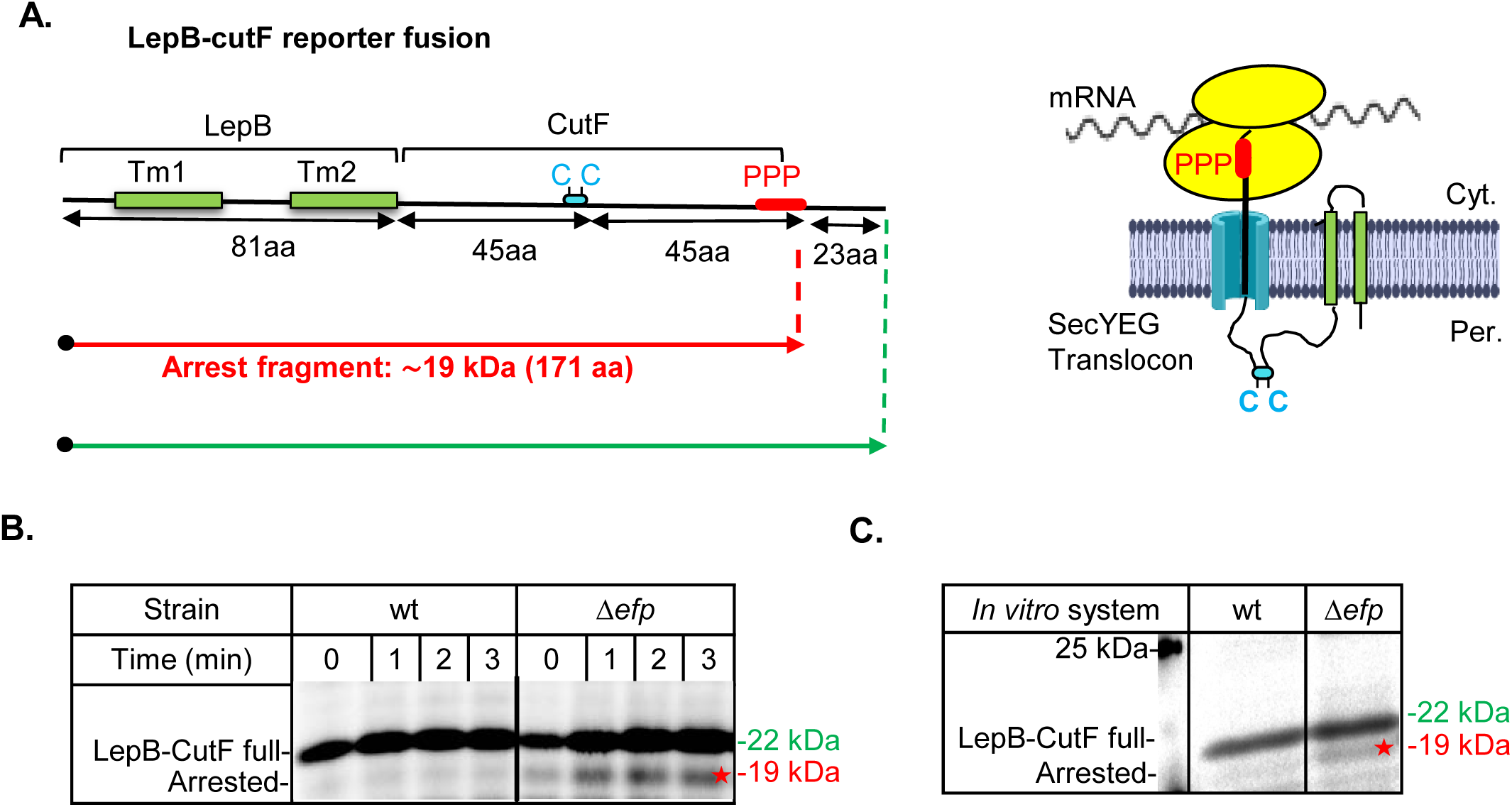
The LepB-cutF reporter fusion is arrested in an Δ*efp* background. **(A)** Cartoon showing the LepB-CutF reporter fusion (left panel). Tm1 and Tm2 refer to the first two transmembrane domains of LepB. The cysteine-containing Cu-binding motif (Cys_41_xxxCys_45_) and the C-terminal proline-rich motif of CutF are indicated, while the upstream-located residues Cys_5_ and Cys_12_ are not shown. The 23 amino acid fragment of LepB after the proline-rich motif was kept to distinguish between the full-length protein and the arrested fragment. The right panel shows a cartoon representation of this construct during cotranslational membrane localization. **(B)** Time-dependent pulse labelling of LepB-CutF in *E. coli* wt and in Δ*efp* strains were performed *in vivo* using ^35^S-labelled methionine/cysteine. After the 0-, 1-, 2-, and 3-min pulses, whole cells were precipitated with trichloroacetic acid (TCA), separated via SDS-PAGE, and analyzed via phosphorimaging. The full-length LepB-CutF and the arrested fragment are indicated. **(C**) An *E. coli in vitro* transcription/translation system isolated from either wt or Δ*efp* cells was employed for the *in vitro* synthesis of LepB-CutF. In vitro synthesized proteins were labelled with ^35^S-methionine/cysteine and analyzed after SDS-PAGE via phospho-imaging.

The presence of the 23 amino acid long C-terminal extension allows to differentiate between the stalled (∼19 kDa) and full-length (∼22 kDa) CutF. *In vivo* radioactive metabolic labeling of the wild type strain showed the 22 kDa LepB-CutF full-length protein, but not its stalled version. In contrast, in the *E. coli* Δ*efp* strain, both the 22 kDa full-length LepB-CutF and the 19 kDa stalled LepB-CutF fragment were observed (**Fig. 4B**). In addition, the 19 kDa stalled LepB-CutF fragment was also observed when CutF was *in vitro* synthesized using a coupled transcription/translation system derived from the Δ*efp* strain (**Fig. 4C**), further confirming the *in vivo* data. These findings further support that CutF being is stalled inside the ribosome tunnel when EF-P is absent and also support a model in which CutO production depends on CutF stalling.

For demonstrating that CutF is indeed stalled inside the ribosomal tunnel (**Fig. 5A**), an *in vitro* site-directed cross-linking approach using the UV-sensitive amino acid derivative para-benzoyl-L-phenylalanine (pBpa) was used (Ryu & Schultz, 2006, Young *et al*., 2010). pBpa was inserted at position 93 of the C-terminus of CutF by an orthogonal tRNA/tRNA synthetase pair during *in vitro* translation. Upon UV-exposure of the *in vitro* reaction mix, two additional UV-dependent bands at approx. 25 kDa were visible, which were not observed for wild type CutF, lacking pBpa **(Fig. 5B)**. The size of these cross-linking products fit in size to cross-links between CutF and ribosomal proteins, such as uL22 (12 kDa) or uL23 (11 kDa). uL23 is located at the ribosomal tunnel exit but also extends via a β-hairpin loop into the ribosomal tunnel, where it acts as a nascent chain sensor (Denks *et al*., 2017, Knüpffer *et al*., 2019). uL22, on the other hand, forms together with uL4 (22 kDa) a narrow constriction within the ribosomal tunnel, that traps many arrest-peptide-containing proteins **(Fig. 5A)** (Harms *et al*., 2001, Seidelt *et al*., 2009).

**Figure 5.**
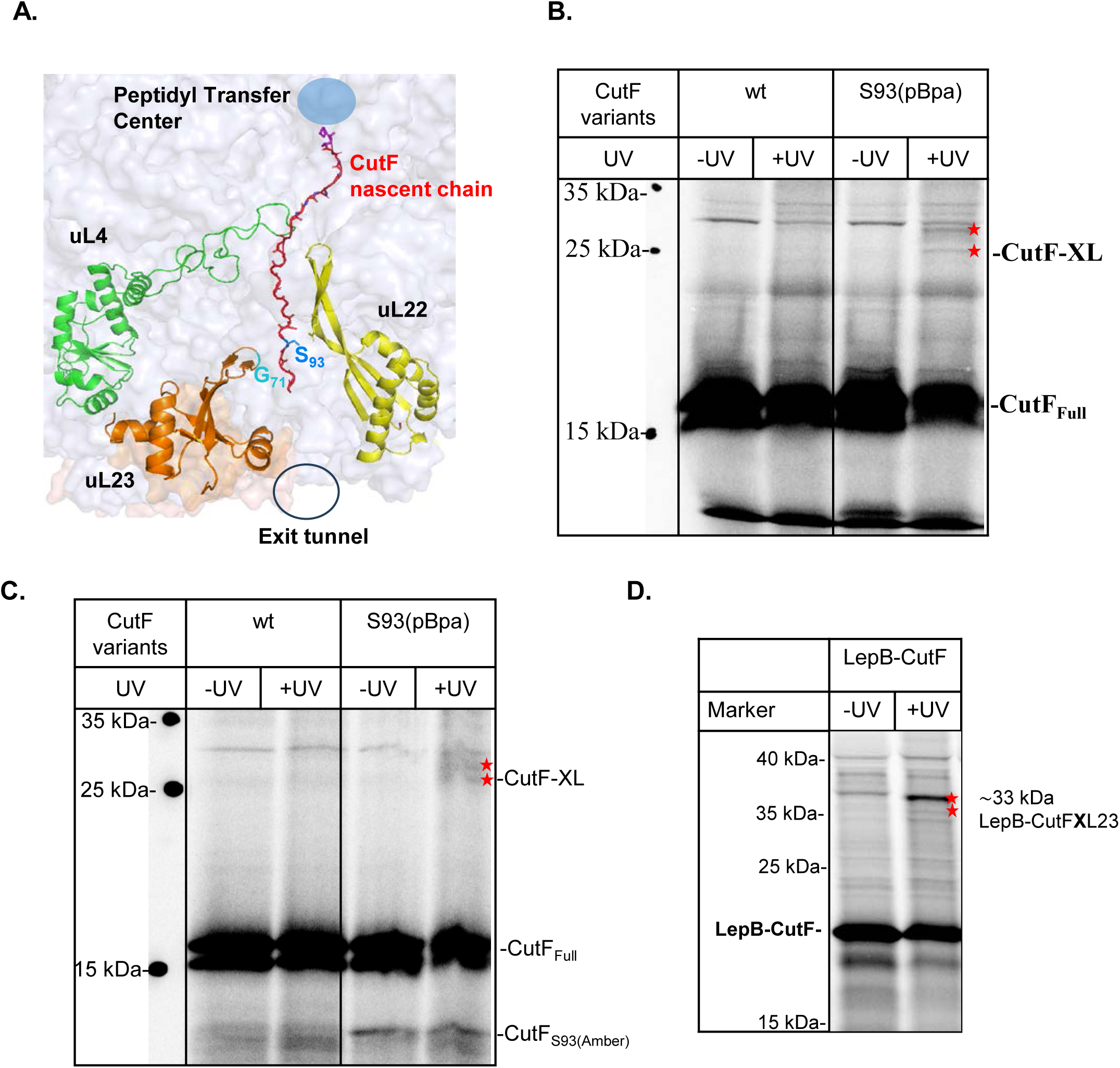
*In vitro* pBpA crosslinking indicates that the C-terminus of CutF is stalled inside of the ribosomal tunnel. **(A)** Cartoon showing the possible orientation of arrested CutF inside the ribosomal tunnel. The PyMOL generated cartoon is based on the structure of stalled SecM inside the ribosome (PDB 3JBV) with CutF replacing SecM. pBpA substitutions in CutF (S_93_, blue, full length numbering) and in the ribosomal protein uL23 (G_71_, cyan) are indicated. **(B)** Wild type CutF or CutF containing pBpa at position 93 were *in vitro* synthesized and during the *in vitro* reaction UV-exposed to induce the cross-linking reaction. As control, the same reaction was kept in the dark (-UV). Samples were then separated on SDS-PAGE and analyzed by phosphorimaging. Possible cross-linking products are indicated (*). **(C)** The ribosomal fraction of the reactions shown in panel (B) were purified by centrifugation through a sucrose cushion and analysed as in panel (**A**). **(D)** *In* vitro synthesis of LepB-CutF was performed on ribosomes that contained a variant of the ribosomal protein uL23 with pBpa inserted at position 71, reaching into the ribosomal peptide tunnel (see panel A). Induction of the cross-link reaction and the analyses were performed as in (**B**).

The potential cross-links between CutF and ribosomal proteins were further validated by isolating ribosome-bound cross-linking products via centrifugation though a sucrose cushion. This procedure let to the enrichment of the two cross-linking bands, which supports a possible crosslink of CutF to ribosomal proteins **(Fig. 5C)**.

We have recently generated functional ribosomes that contain a uL23 variant with pBpa inserted at position G71 at the tip of the β-hairpin loop (Denks *et al*., 2017, Knüpffer *et al*., 2019) **(Fig. 5A)**. When these ribosomes were used for *in vitro* synthesis of LepB-CutF, two cross-linked bands of approx. 33 kDa and different intensities were observed that fit in size to LepB-CutF-uL23 cross-linking products **(Fig. 5D)**. The observation that cross-linking products migrate in several bands is frequently observed with pBpa cross-linking and reflects different conformations of the cross-linking product (Kuhn *et al*., 2015, Kuhn *et al*., 2013, Mori & Ito, 2006, Das & Oliver, 2011). The overall data thus support that CutF-stalling inside the ribosome is a prerequisite for CutO production. However, it remains to be analyzed how Cu binding in the periplasm can induce ribosomal stalling of CutF by alleviating the function of EF-P.

## Discussion

Bacteria depend on sophisticated mechanisms that allow them to process environmental signals and to integrate them into cellular responses. This is not only important for metabolic adaptation to different nutrient availability, but also for evading or detoxifying potentially damaging substances, such as heavy metals.

Here, we have identified a remarkable sensing mechanism that allows *R. capsulatus* to sense the periplasmic Cu concentration via the nascent CutF protein and to use this information to regulate the translation initiation of the detoxifying copper oxidase CutO. Periplasmic Cu sensing is well established for two-component systems, such as the CusRS system of *E. coli* (Munson *et al*., 2000, Andrei *et al*., 2020). In this system, the histidine kinase CusS senses the periplasmic Cu concentration and phosphorylates the response regulator CusR, which in turn transcriptionally activates the CusABC Cu export system (Long *et al*., 2010, Mealman *et al*., 2012, Su *et al*., 2011). The CutFO system described in our study uses an entirely different mechanism that is based on Cu-induced ribosomal stalling of CutF. This in turn allows the ribosome via its mRNA helicase activity (Takyar *et al*., 2005) to melt an mRNA stem loop that shields the *cutO* ribosome-binding site, allowing for translation initiation and production of CutO **(Fig. 6 A)**. Although regulating protein synthesis via translational stalling has been demonstrated before and is well characterized for the regulated production of SecA (Ito *et al*., 2018), SecDF (Ishii *et al*., 2015) or TnaAB (Seidelt *et al*., 2009, Martínez *et al*., 2014), the CutFO system is to our knowledge the first system that senses extra-cytoplasmic ligands and converts this information into transmembrane stalling.

**Figure 6.**
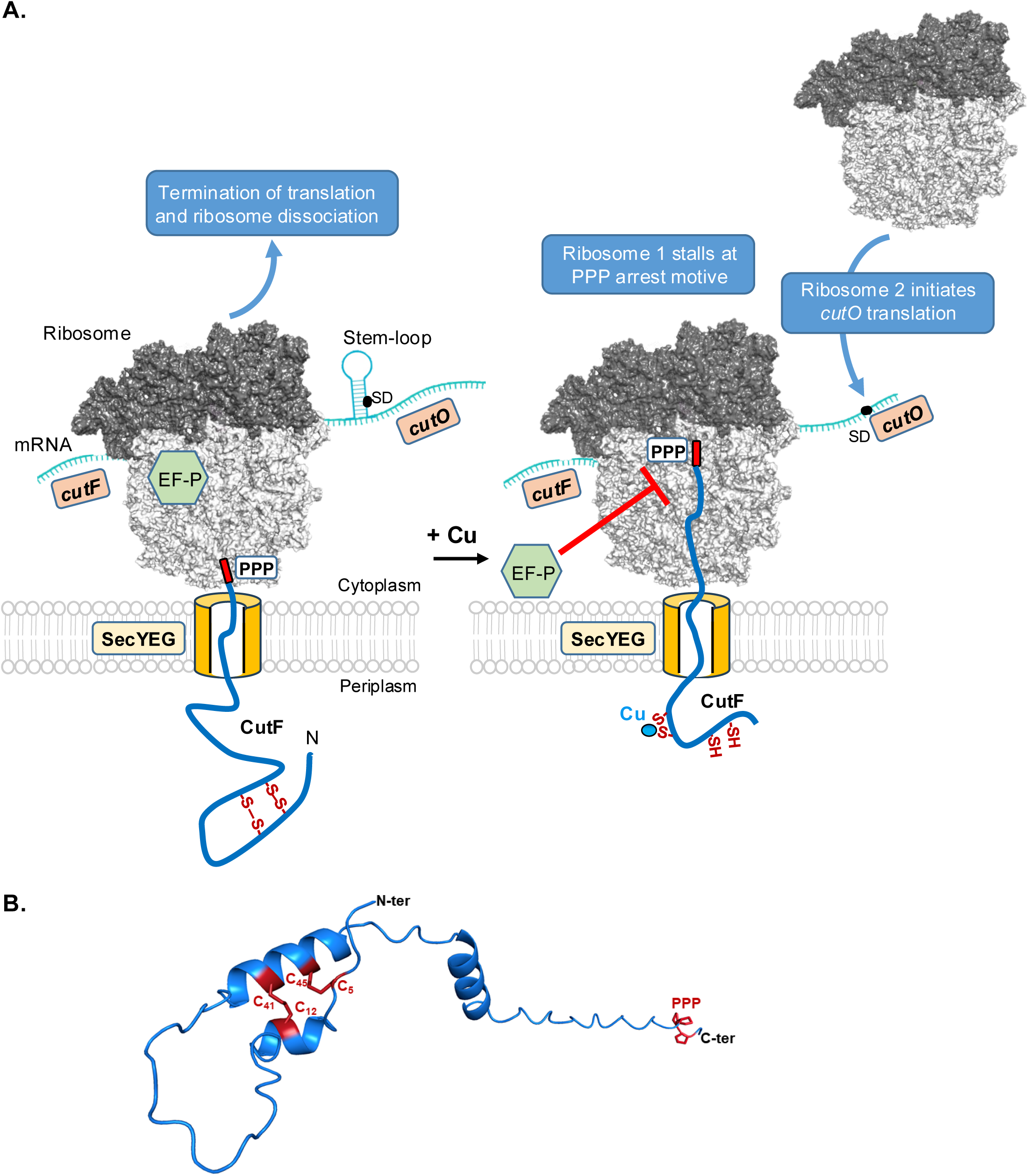
Model for Cu-dependent CutO production via ribosomal stalling of CutF. **(A)** CutF is co-translationally targeted to the SecYEG translocon and secreted into the periplasm. In the absence of Cu (left panel), the cysteine residues within the Cu-binding C_41_XXXC_45_ motif, along with two additional upstream-located cysteine residues (Cys_5_ and Cys_12_), likely form disulfide bridges in the oxidizing environment of the periplasm. The formation of this secondary structure, together with the activity of EF-P and the pulling force of the SecYEG translocon is sufficient to relief stalling at CutF’s C-terminal PPP motif and allows complete synthesis of CutF. Consequently, the downstream stem-loop structure remains intact, shielding the Shine-Dalgarno (SD) sequence of *cutO* and preventing its translation. In the presence of Cu (right panel), the nascent CutF protein binds Cu via its periplasmic CXXXC motif. The inability to form disulfide-bridges leads to the arrest of the PPP motif within the ribosomal tunnel, which cannot be rescued by EF-P. This transient stalling provides sufficient time for the helicase activity of the ribosome to unfold the stem-loop structure, exposing the SD sequence and enabling the initiation of *cutO* translation. Stalling in the presence of Cu might be further enhanced by Cu-induced impairment of EF-P modifying enzymes or of the protein secretion machinery. Currently, we can not differentiate whether *cutO* translation is initiated by a second ribosome, as depicted in the cartoon, or by the stalled ribosome, after stalling is relieved. The ribosome structure was generated by using PyMOL based on the structure of the ribosome with EF-P (PDB 6ENJ) (Huter *et al*., 2017). **(B)** Structural prediction of mature CutF generated by AlphaFold3. The amino acid sequence corresponds to the mature form of CutF following signal peptide processing. Cysteine residues, predicted disulfide bridges, and the C-terminal PPP repeat are highlighted in dark red.

CutF stalling strictly depends on its C-terminal PPP-motif. Prolines are generally poor substrates for peptide-bond formation, because they destabilize the P-site tRNA (Hummels & Kearns, 2020, Doerfel & Rodnina, 2013) and the PPP-motif is a particularly strong staller (Woolstenhulme *et al*., 2013, Peil *et al*., 2013). However, ribosomal stalling at consecutive proline residues is usually prevented by EF-P (Ude *et al*., 2013, Doerfel *et al*., 2013), which stabilizes the P-site tRNA and accelerates peptide-bond formation between proline residues (Huter *et al*., 2017). Our data show that Cu-binding to the periplasmically exposed Cu-binding motif of CutF overrides the function of EF-P and causes ribosomal stalling. On the other hand, when EF-P is deleted CutF stalling and CutO production becomes Cu-independent. Intriguingly, the signal sequence of CutF is required for stalling and CutO production, which indicates that stalling is linked to co-translational secretion of CutF (Öztürk *et al*., 2023). Due to the presence of Cu chaperones, the intracellular concentration of free Cu is very low, even when cells are grown on Cu-supplemented media (Andrei *et al*., 2020, Utz *et al*., 2019, Capdevila *et al*., 2024). Thus, one possibility is that the cytosolic Cu concentration is not sufficient to cause stalling. In addition, although the absence of EF-P causes stalling and CutO production, this is significantly impaired when the signal sequence is deleted (**Fig. S3**). These findings suggest that the signal sequence or co-translational secretion plays a particular role in CutF stalling (**Fig. 6A**). For SecM it was shown that stalling is released by a mechanical pulling force caused by the interaction of the N-terminal signal sequence of SecM with SecA (Butkus *et al*., 2003, Ito & Chiba, 2013). CutF is co-translationally targeted to the SecYEG translocon by the signal recognition particle (SRP) (Öztürk *et al*., 2023, Steinberg *et al*., 2018, Oswald *et al*., 2021). However, this does not exclude that after the initial SRP-dependent targeting, the ATPase SecA might be involved in translocating CutF across the SecYEG translocon. Such a coordination of SRP-dependent targeting with SecA-dependent translocation is required for the transport of many bacterial proteins (Neumann-Haefelin *et al*., 2000, Scotti *et al*., 1999). Therefore, it is possible that stalling of CutF in the absence of Cu is relieved by the pulling force of SecA and/or the SecYEG-translocon (Niesen *et al*., 2018, Goldman *et al*., 2015), combined with the activity of EF-P. The situation is different in the presence of Cu or when EF-P is deleted, under these conditions stalling occurs demonstrating that the pulling force is not sufficient. Molecular dynamics simulations demonstrate that secondary structures modulate the force required for the relief of SecM stalling (Gersteuer *et al*., 2024). Thus, a change in the secondary structure of the nascent CutF upon Cu binding could increase the force required for stalling relief.

CutF contains two cysteine residues within its periplasmic Cu binding motif (Cys_41_ & Cys_45_) and two additional cysteine residues upstream within the mature domain (Cys_5_ & Cys_12_, mature CutF numbering). Alphafold predictions suggest the presence of two possible disulfide bridges (Cys_5_-Cys_45_ and Cys_12_-Cys_41_), which can contribute to the relief of stalling (Elfageih *et al*., 2020) (**Fig. 6 B**). The pulling force generated by the formation of the two disulfide bridges would not be possible when Cu occupies the Cu-binding motif and the associated secondary structure changes could even further increase the force required for stalling relief.

The mechanism of how the addition of Cu overrides the function of EF-P is currently unknown. *In vivo*, the activity of EF-P is strongly enhanced by lysinylation of a conserved lysine residue (lysine 34 based on *E. coli* numbering) (Doerfel & Rodnina, 2013), which is catalyzed in *E. coli* and many other bacteria by the three proteins, YjeK, YjeA and YfcM (also called EpmA, EpmB and EpmC) (Peil *et al*., 2012). YfcM hydroxylates lysine 34 and this hydroxylation appears to stabilize the tRNA conformation in the P-site (Kobayashi *et al*., 2014). The activity of YfcM depends on a 2-His-1-carboxylate motif that coordinates Fe(II) and which is typical for non-heme iron enzymes (Que, 2000, Kobayashi *et al*., 2014). Intriguingly, Cu toxicity is primarily caused by inhibition of FeS cluster biogenesis and by displacing other metals in already assembled enzymes (Capdevila *et al*., 2024). Thus, one possibility is that the addition of Cu increases the cytosolic Cu concentration to a level that inactivates YfcM and in turn impairs the activity of EF-P; but this needs to be further analyzed. It is important to emphasize that even transient stalling would be enough for melting such a stem-loop (Ito & Chiba, 2013) and for this a partial reduction of EF-P activity would likely suffice.

Although the DUF2946 family of proteins is widely distributed in bacteria, the absence of EF-P shows different effects on the stalling of the two best-studied prototypes, CutF and CruR. While stalling of CruR in *Bordetella pertussis* was not affected in a Δ*efp* strain (Roy *et al*., 2022), our data clearly show that EF-P is involved in relieving the stalling of CutF. A possible explanation comes from the observation that ribosome stalling at polyprolines sites can be rescued not only by EF-P but also by other enzymes, such as EfpL, Uup or YfmR (Sieber *et al*., 2024).

In conclusion, our data reveal a new post-transcriptional strategy that bacteria use to cope with the essential and yet toxic heavy metal Cu. Considering the wide distribution of DUF2946 proteins within bacteria, this strategy is most likely used by many bacteria for regulating their metal homeostasis.

## Acknowledgements

This work was supported by grants of the Deutsche Forschungsgemeinschaft to HGK (SFB1381, Project-ID 403222702, and RTG 2202, Project-ID 278002225). YÖ acknowledges support by the Philipp Schwartz Initiative of the Alexander von Humboldt Foundation and by the RTG 2202.

## Author Contributions

Conceptualization: HGK, FD; Investigation: YÖ, KWS, PE; Visualization: YÖ, FD, HGK; Funding acquisition: FD, HGK; Supervision: YÖ, HGK, FD; Writing: YÖ, FD, HGK. All authors have read the manuscript.

## Supplemental information

Document S1. Figure S1-S3 and Tables S1 and S2

