## Supplemental information for "Copper-controlled gene expression via transmembrane-induced ribosomal stalling"

**A.**

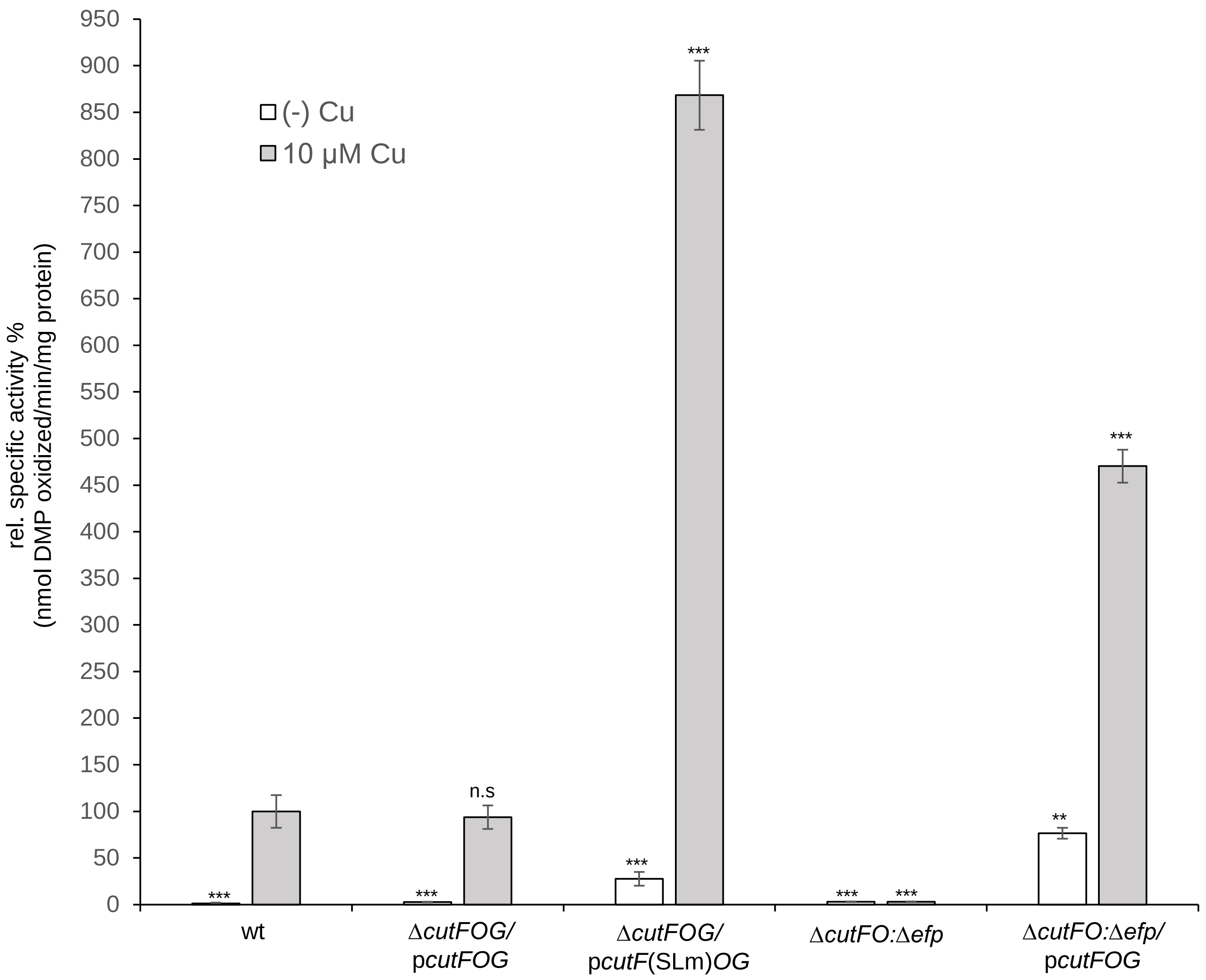


**B.**

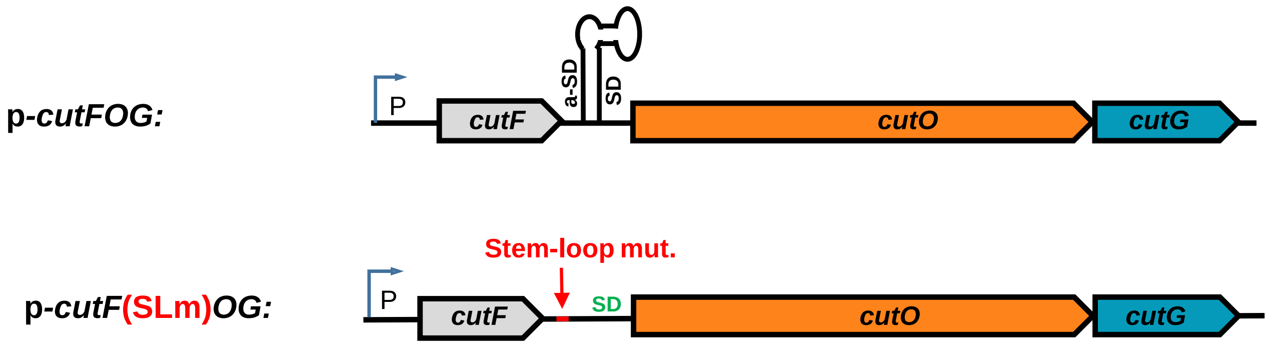


(Öztürk et al., Figure S1)

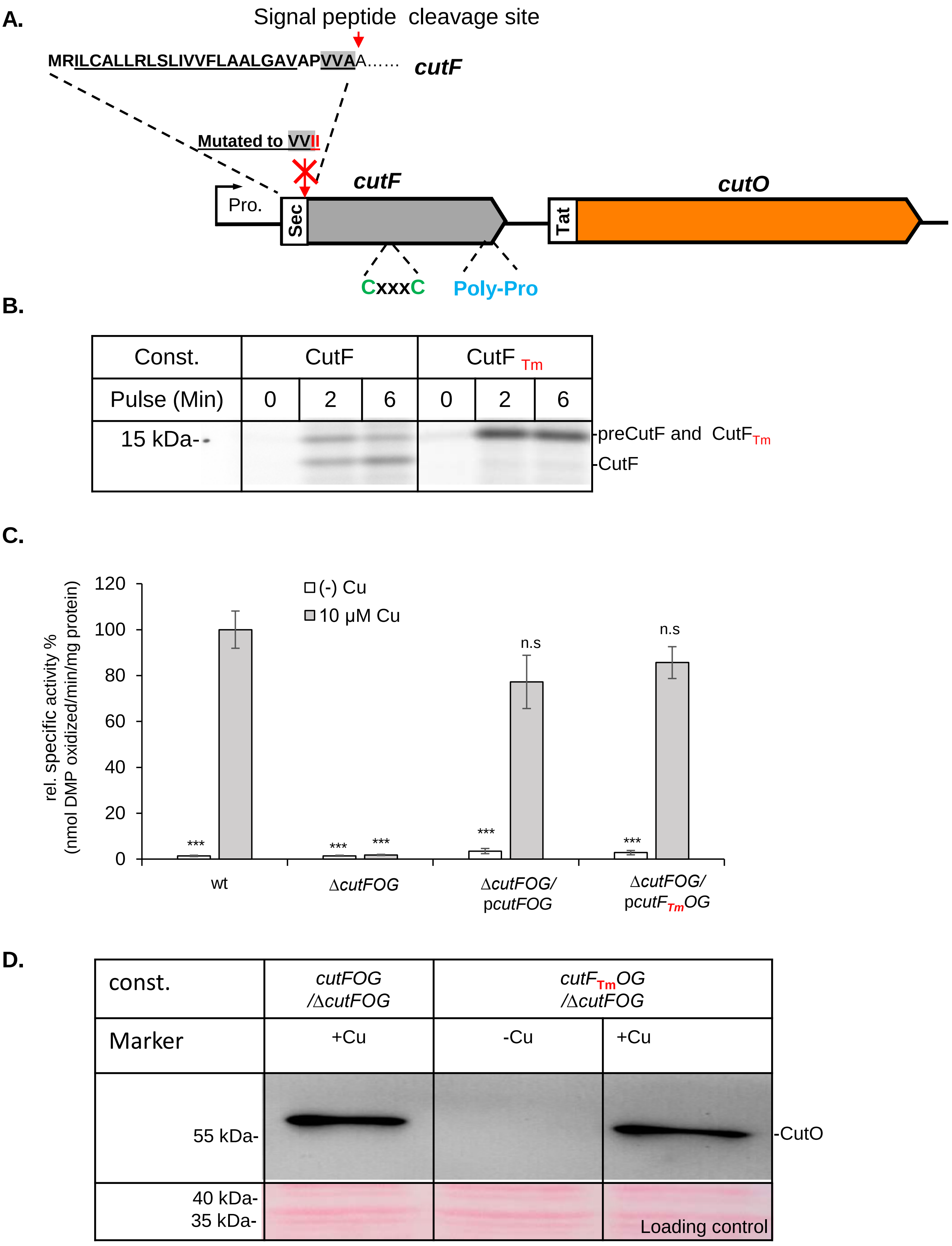


(Öztürk et al., Figure S2)

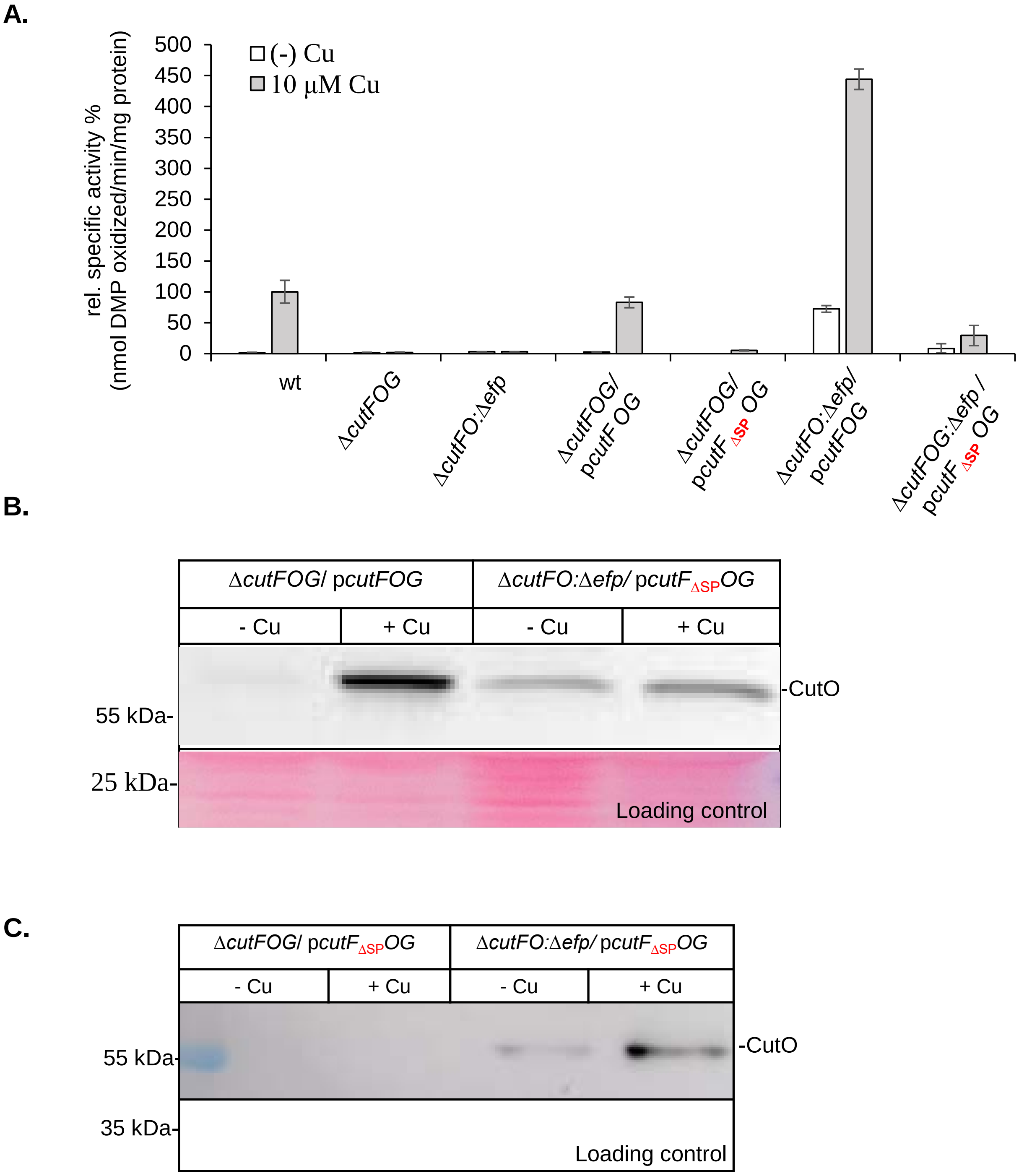


(Öztürk et al., Figure S3)

### Legends to Supplementary Figures:

**Figure S1: CutO activity in different *R. capsulatus* strains (A)** CutO activity of *R. capsulatus* strains grown on MPYE medium supplemented without or with 10 mM $CuSO_4$. p*cutFOG* encodes the entire *cutFOG* operon and *pcutF*(SLm)*OG* the entire operon with a mutation within the stem-loop separating *cutF* and *cutO.* Periplasmic fractions were isolated from the indicated strains and 50 mg periplasmic soluble protein were used for CutO activity as described in the legend to Fig. 1B. (**B**) Cartoon showing the genetic organization of the *cutFOG* operon and the presence of the stem-loop.

**Figure S2. A membrane-bound CutF still functions as Cu-dependent regulator of CutO synthesis. (A)** Cartoon showing the substitution of the signal peptide cleavage site VVAA to VVII, generating the membrane-bound CutF variant $CutF_{TM}$. **(B)** *In vivo* pulse labeling of CutF and $CutF_{Tm}$ for 0-, 2-, and 6-min. The samples were precipitated and analyzed via phosphorimaging as described in Fig. 4B. **(C)** The CutO activity was determined by using 50 mg periplasmic soluble total proteins as described in the legend of Fig. 1B. **(D)** Immunoblot analyses of the strains harboring $CutF_{Tm}$ variant. 100 mg of periplasmic soluble protein were separated on 15% SDS PAGE and a-Flag antibodies were used for the detection of CutO as described in Materials and Methods. The same membrane was stained with Ponceau solution as loading control.

**Figure S3. The signal sequence of CutF is required for regulating CutO production. (A)** CutO activity of *R. capsulatus* strains carrying a CutF variant that lacks the signal sequence ($CutF_{\Delta SP}$) in the presence and absence of EF-P**.** 50 mg periplasmic soluble total proteins as described in the legend to Fig. 1B. **(B) and (C)** 100 mg of the periplasmic fractions of the indicated strains, grown in the absence or presence of 10 µM Cu were used for immunoblot analyses. After separating on 15% SDS PAGE and transfer to a nitrocellulose membrane, a-Flag antibodies were used for the detection of CutO as described in Materials and Methods. The same membrane was stained with Ponceau solution for loading control.

**Table S1.** Strains and plasmids used in this work.

| Strain or plasmid | Description | Phenotype | Reference |
|---|---|---|---|
| ***Strains***<br><br>***E. coli*** | | | |
| HB101 | $F^-$ Δ(*gpt-proA*)62 *leuB6 supE44 ara*-14 *galK2 lacY*1<br>Δ(*mcrC-mrr*) *rpsL20* ($Str^R$) *xyl-5 mtl-1recA*13 | $Str^r$ | [1] |
| NEB 5-α | *fhuA2 D(argF-lacZ)U169 phoA glnV44 f80D(lacZ)M15 gyrA96 recA1*<br>*relA1 endA1 thi-1 hsdR17* | | [2] |
| MC4100 | *Wild type strain* | | [3] |
| BW25113 | *Δ(araD-araB)567, ΔlacZ4787*(::rrnB-3), *λ*⁻, *rph-1, Δ(rhaD-rhaB)568, hsdR514* | | [4] |
| JW4107(Δ*efp*) | *Δ(araD-araB)567, ΔlacZ4787*(::rrnB-3), *λ*⁻, *rph-1, Δ(rhaD-rhaB)568, Δefp-772::kan, hsdR514* | $Kan^r$ | [5] |
| C43 (DE3) | *F*⁻*, ompT, gal, dcm, hsdS*$_B$*(r*$_B^-$ *m*$_B^-$*)(DE3)* | | [6] |
| <br>***R. capsulatus*** | | | |
| [a]MT1131 | *crtD121* | Wt ($Rif^r$) | [7] |
| Y262 | GTA overproducer | | [8] |
| Δ*cutF*<br>(YO-ΔcutF) | Δ( *cutF*(*rcc02111*)) seamless in frame deletion | | [9] |
| Δ*cutFOG*<br>(YO-ΔcutFOG) | Δ ( *rcc02111*), Δ(*rcc0209::Gm*), Δ(*cutO::Kan*) | $Kan^r$, $Gm^r$ | [10] |
| Δ*efp* | Δ(*efp::Gm*) | | This work |
| Δ*cutFO& Δefp* | Δ ( *rcc02111*), Δ(*cutO::Kan*), Δ(*efp::Gm*) | $Kan^r$, $Gm^r$ | This work |
| <br>***Plasmids*** | | | |
| pRK2013 | Conjugation helper | $Kan^r$ | [11] |
| pRK415 | Broad host-range vector | $Tet^r$ | [11] |
| pRS1 | pPET19 based containing pBR322 ori, Rop, T7RNAP under EM7 promoter, LacI | $Amp^r$ | [12] |

| | | | |
|---|---|---|---|
| p*cutFOG* | *cutF*$_{\text{N-ter Flag}}$, *cutO*$_{\text{Flag}}$ and *cutG*$_{\text{MycHis}}$ with promoter and terminator regions of *cutFOG* operon on pRK415 | $Tet^r$ | [10] |
| pCHB::Gm | | $Gm^r$, $Tet^r$ | [9] |
| pEVOL | tRNA synthetase/tRNA pair for the in vivo incorporation of a photocrosslinker, p-azido-l-phenylalanine (pBpA), into proteins in *E. coli* response to the amber codon, TAG | $Chm^r$ | [13] |
| p*cutF*$_{\text{C-A}}$*OG* | substitution of conserved Cys to Ala of CutF ($C_{69}XXXC_{73}$ to $A_{69}XXXA_{73}$) on pcutFOG | $Tet^r$ | [10] |
| p*cutF*$_{\text{SLm}}$*OG* | CTTC to AAAA mutation on anti-SD of Steem-loop (SLm) between the *cutF-CutO* intergenic region on p*cutFOG*. | $Tet^r$ | [14] |
| p*cutF*$_{\Delta\text{SP}}$ *OG* | truncation of Sec signal peptide of CutF on p*cutFOG*. | $Tet^r$ | [14] |
| p*cutF*$_{\text{TM}}$*OG* | Signal peptide cleave site $VA_{28}A_{29}$ to $VI_{28}I_{29}$ substitution on p*cutFOG* to obtain transmemrane attached CutF | $Tet^r$ | This work |
| p*cutF*$_{\text{PAP}}$*OG* | $PP_{115}P$ to $PA_{115}P$ substitution of CutF on p*cutFOG*. | $Tet^r$ | This work |
| p*efp* | *efp* gene with promoter and terminator regions on pRK415 | $Tet^r$ | This work |
| p*efp::Gm* | Δ(*efp::Gm*) on pRK415. | $Tet^r$, $Gm^r$ | This work |
| pRS-CutF | *cutF*$_{\text{N-Flag}}$ ORF cloned to pRS1 | $Amp^r$ | [10] |
| pRS-CutF$_{\text{S93pBpA}}$ | S93Amber SC substitution of CutF$_{\text{N-Flag}}$ on pRS1 | $Amp^r$ | This work |
| pRSLebB-CutF | ORF of *cutF* (excluding SP) fused to LepB after 2$^{nd}$ transmembrane helix | $Amp^r$ | [14] |

[a]*R.capsulatus* strain MT1131 is derived from SB1003 in multiple steps, as described in[7]: first, a Ps-deficient mutant (TL1) was obtained using tetracycline suicide, then its *crtD* derivative was constructed by GTA cross to yield MT113, and then its Ps-proficient derivative was obtained via a second GTA cross.

**Table S2.** Primers used in this work.

| | |
|---|---|
| cutF(EP)-F | 5-agtgaattcgagctcggtacatcctgcgcctgaaaggccag-3 |
| cutTer-R | 5-tgcatgcctgcaggtcgactaatagtctctttggcctgctgccg-3 |
| cutFstop&noSS-R | 5-gggaggagcatggcacggggcgggg-3 |
| EFP-F | 5- agtgaattcgagctcggtaccggcgcagtcctgcatcagcacgg-3 |
| EFP-R | 5-tgcatgcctgcaggtcgactttgcatgtcggcccgcaacggcggtc-3 |
| EFP-Fr1-R | 5-cagatcgggtcgacaagcttctgcgtcaccgagttgcccttgccc-3 |
| EFP-Fr2-F | 5-ggggatccactagttctagacgatcgaactgccgcaaaaggtcacc-3 |
| Gm-F | 5-aagcttgtcgacccgatctgagc-3 |
| Gm-R | 5-tctagaactagtggatcccccg-3 |
| pRS1-F | 5-gcggccgcgagctcgtcgacctcgagtagc-3 |
| CutF(P115A)-F | 5-gctgctttcccccgaaccggaggggccggcgccccgcctctgatcttcgc-3 |
| CutF-CterM-Rv | 5-ttcgggggaaagcagcgccagccgc-3 |
| CutF(SPm)-F | 5-ggtcatcatcgactacaaggacgacgatgacaagggcggctcggcgccggaggcctgtc-3 |
| CutF(SPm)-R | 5-ccttgtagtcgatgatgaccacgggcgcaaccgcgccaagc-3 |
| pRS1-R | 5-catggggtatatctccttcttaaagttaaacaaaattatttc-3 |
| pRS1cutF-R | 5-gtcgacgagctcgcggccgctcagaggcggggcggcggcccctcc-3 |
| pRScutF(noSS)-F | 5-gcggactacaaggacgacgatgacaagg-3 |
| CutF(SPm+Flag)-F | 5-atcatcgactacaaggacgacgatgacaagggcggc-3 |
| CutF(SP-mut)-R | 5-gaccacgggcgcaaccgcgccaagc-3 |
| cutFnoSS-F | 5-tgccatgctcctcccatggcggactacaaggacgacgatgacaagggcggc-3 |
| pRScutF(P-A)O-F | 5-gccggcgccccgcctctgatcttcgccacgttcccattcttgaacc-3 |
| pRScutF(P-A)O-R | 5-atcagaggcggggcgccggcccctccggttc-3 |
| cutO-SL-F | 5-gccgccgccccgcctctgataaaagccacgttcccattcttgaacc-3 |
| cutO-SL-R | 5-atcagaggcggggcggcggcccctcc-3 |
| cutF(CtoA)-F | 5-gcgctgcagcatgcgctgggcgcctcgctgacgcccg-3 |
| cutF(CtoA)-R | 5-gccttccggcgcgaaggtggtggtcttgg-3 |
| CutF(S93TAG)-F | 5-agcgccttgttccgcctgcggttc-3 |
| CutF(AmberTGA)-R | 5-agatgcgctctgcccttgcg-3 |
